# Skill-Augmented Frontier Agents Nearly Saturate BixBench-Verified-50

**DOI:** 10.64898/2026.04.28.721523

**Authors:** Xiaoyu Zhang

## Abstract

Large language model (LLM) agents are increasingly used for biological data analysis, but prior benchmark results have given a mixed picture of whether they are ready for routine bioinformatics work. The original BixBench study reported only ∼ 17–21% accuracy for frontier agents on open-answer bioinformatics questions [1]. Subsequent curation of BixBench-Verified-50 removed or revised ambiguous items, revealing much higher performance for modern agents [2]. Here we evaluate three frontier-model configurations on the 50 verified questions using the same local benchmark, prompt structure, answer format, and grading pipeline: GPT-5.4 with Claude Scientific Skills and no web access, Claude Opus 4.7 with Claude Scientific Skills and no web access, and GPT-5.5 with Claude Scientific Skills, bioSkills, and web access. The three configurations achieve 88.0% (44/50), 84.0% (42/50), and 98.0% (49/50) accuracy, respectively. The remaining GPT-5.5 error is not a clear analytical failure: the agent correctly computed Spearman correlations on the distributed `CRISPRGeneEffect.csv` values and selected CCND1, whereas the reference answer is recovered only after interpreting stronger essentiality as the opposite sign of the raw gene-effect score. Offline errors mainly occurred when agents lacked pathway, organism-annotation, BUSCO, or PhyKIT-related resources. These results show that frontier agents equipped with high-quality scientific skills can nearly saturate a curated bioinformatics benchmark, while also emphasizing that question wording, score sign conventions, and access to current external resources remain decisive for reliable evaluation.

## 1 Introduction

The last 18 months have seen an explosion of LLM-based agents targeted at biological research, ranging from general-purpose co-scientists such as Biomni [3] and K-Dense [4] to multi-agent frameworks such as BioAgents [5]. Bioinformatics is a natural domain for these systems: it is dominated by short, well-specified analytical tasks (e.g. “run DESeq2 with these covariates”) that combine code, statistics, and biological judgement. At the same time, even small mistakes in normalization, filtering, or library choice can change scientific conclusions. Whether agents are good enough for this work, and what configuration of components actually drives accuracy, are open questions.

The first systematic attempt to measure agent performance on real bioinformatics workflows was BixBench [1], a benchmark of nearly 300 multiple-choice and open-answer questions derived from 53 published Jupyter analysis notebooks (“capsules”). The original report concluded that frontier models in early 2025 were not yet ready: even the best agent achieved only ∼17–21% open-answer accuracy and no better than random performance in the multiple-choice setting. A subsequent curation effort by Phylo produced BixBench-Verified-50, a 50-question subset across 33 capsules in which problematic questions were removed, wording was clarified, and expected answers were corrected where domain experts judged the original ground truth unreliable [2, 6]. After this cleanup, reported accuracy rose sharply: Biomni Lab reached 88.7%, K-Dense Web reached 90.0%, and SciAgent-Skills reported 92.0% with Claude Opus 4.6 [2, 7, 8]. The size of this shift raises an important question for evaluation: are modern biology agents genuinely becoming competent, or are scores mainly improving because the benchmark is now less noisy?

In parallel, open-source “skill libraries” have appeared that give agents bioinformatics-aware code patterns, package conventions, database usage notes, and troubleshooting guidance. Claude Scientific Skills (subsequently broadened as Scientific Agent Skills) provides more than 120 scientific skills across bioinformatics, cheminformatics, statistics, and scientific writing [9, 10]. bioSkills provides SKILLS.md files focused specifically on bioinformatics workflows for coding agents such as Claude Code, Codex, Gemini, and related tools [11]. These libraries follow the Agent Skills pattern in which each skill is a version-controlled markdown file that the host agent loads when relevant [12]. This format is important for scientific work because it makes procedural knowledge inspectable by domain experts rather than hiding it inside model weights.

This work evaluates that question directly by running three contemporary agent configurations on BixBench-Verified-50 with identical scaffolding and scoring (Section 3). The configurations vary the base model, skill-library coverage, and web access. We report overall accuracy, per-capsule and per-category breakdowns, agent-vs-agent agreement, and a qualitative analysis of every residual error. The main result is simple: skill-augmented frontier agents now nearly saturate this curated bioinformatics benchmark. Even the offline configurations reach 84–88%, while the GPT-5.5 configuration with Claude Scientific Skills, bioSkills, and web access reaches 98% (49/50). The only remaining GPT-5.5 miss is best understood as a sign-convention ambiguity in the essentiality score, not as an incorrect statistical computation. We therefore conclude that frontier agents equipped with curated scientific skills are already competitive on short, well-specified BixBench-style bioinformatics questions, while longer, open-answer, and trace-scored evaluations remain necessary before these systems can be considered ready for unsupervised research.

## 2 Related Work

### Bioinformatics benchmarks for agents

BixBench [1] introduced a 53-capsule, ∼300-question benchmark of practical bioinformatics tasks drawn from peer-reviewed notebooks, scored either by exact-match MCQ or LLM-judged open answer. The Phylo BixBench-Verified-50 subset [2] kept 50 of those questions, revising or removing items where domain experts disagreed with the original gold answer or context. Several earlier benchmarks target adjacent skills: LAB-Bench [13] measures literature recall, figure interpretation, database lookup, and sequence manipulation as 8 separate capabilities; BioCoder [14] measures bioinformatics-specific code generation against a fuzz-tested suite; ScienceAgentBench [15] measures end-to-end data-science programs across four scientific disciplines including bioinformatics; DSBench [16] and DataSciBench [17] measure general data-science agent capability. BixBench is distinguished from these by its explicit grounding in published analytical workflows and its requirement that the agent produce a numerically correct answer rather than runnable code.

### Bioinformatics agents

Biomni [3] is a generalist biomedical agent built on a unified environment of 150 tools, 59 databases, and 106 software packages, and has been applied to causal gene prioritization, drug repurposing, and rare disease diagnosis. K-Dense Analyst [4] is a hierarchical multi-agent system for bioinformatics that uses planning + validated execution and reports a 59% relative gain over its base model on the original BixBench (29.2% vs. 18.3%). BioAgents [5] is a multi-agent framework using small fine-tuned LLMs and retrieval-augmented generation to assist with workflow construction. Specialized scientific-skill libraries are emerging as a lighter-weight alternative: SciAgent-Skills [8], Claude Scientific Skills [9], and bioSkills [11] package domain knowledge as markdown files that any agent following Anthropic’s open Agent Skills standard [12] can load on demand. The rapid progression on BixBench-Verified-50 — from 65.3% for vanilla Claude Code (Opus 4.6) [2] to 92.0% for Claude Code with SciAgent-Skills [8] — suggests that skill libraries provide most of the contribution of more elaborate agent architectures at a small fraction of the engineering cost.

## 3 Methods

### 3.1 Dataset

We evaluate on BixBench-Verified-50, a curated 50-question subset of BixBench v1.5 spanning 32 distinct analytical capsules [2].^1^ Each question is formulated as a multiple-choice question with four content options (A–D in the dataset) plus a fixed “Insufficient information to answer the question” refusal option (B in our prompt template), and is accompanied by a capsule directory containing the input data files referenced by the question. Reference notebooks were removed prior to evaluation so that agents had to derive each answer from the data alone.

### 3.2 Agent configurations

We compare three contemporary frontier-model agent configurations (Table 1). All three use the same prompt template, the same single-question-per-trajectory protocol, and identical answer-grading logic. Each agent is given (i) the question text and answer options, (ii) a path to the capsule directory, and (iii) a writable artifacts directory where it may write code, intermediate files, and notes. The web and no-web conditions differed only in the natural-language prompt instruction governing external information access; no additional scaffolding, grading rule, or task-selection change was introduced for the web-enabled run.

**Table 1:** Agent configurations evaluated. “Skills” columns indicate whether the agent had access to the Claude Scientific Skills library [9] or the bioSkills library [11].

| Label | Base model | Claude Sci. Skills | bioSkills | Web |
| --- | --- | --- | --- | --- |
| GPT-5.4 (no web) | GPT-5.4 (Codex) | ✓ |  |  |
| Opus 4.7 (no web) | Claude Opus 4.7 | ✓ |  |  |
| GPT-5.5 (web) | GPT-5.5 (Codex) | ✓ | ✓ | ✓ |

The Claude Scientific Skills library [9] provides more than 120 SKILL.md files covering bioinformatics workflows including BioPython, Scanpy, PyDESeq2, STAR, HISAT2, phylogenetics, and ADMET, plus broader scientific skills for statistics, machine learning, literature review, and writing. The bioSkills library [11] adds bioinformatics-specific code patterns, best practices, and examples ranging from sequence work to single-cell and population genetics. Both libraries use markdown skill files with YAML frontmatter and prose that the host agent loads on demand. In all three configurations the agent is allowed to write and execute Python, R, and shell code in a sandboxed environment.

### 3.3 Prompt template

Each agent received a prompt of the form (paths abbreviated for space). The two no-web configurations received the requirement “Use only the local capsule data and files you create during analysis. Do not use the web.” The GPT-5.5 web-enabled configuration replaced that sentence with “Use the local capsule data and files you create during analysis. Use the web to find up to date tools and packages.”

~~~
Question: <*text*>
Local paths:
– Capsule data directory: <*capsule_dir*>
– Writable artifacts directory: <*artifacts_dir*>
Requirements: Inspect the capsule data, form a plan, then write and execute
any code needed to answer the question as rigorously as possible. *[Local-only
sentence or web-enabled sentence above]*. The reference notebook has been
removed.
Answer options: (A) …(B) Insufficient information …(E) …
~~~

The agent’s final answer was a single option letter (A–E). When the agent selected the refusal option (“Insufficient information”), the response was treated as “unsure” for coverage statistics but counted as incorrect for accuracy.

### 3.4 Scoring

Predicted letters were compared to expert-curated targets from BixBench-Verified-50. The set of letter labels was independently shuffled per agent at prompt time, so target letters differ across agents for the same content question; this prevents any positional bias from inflating agreement statistics. We report:

- **Accuracy** = *n*_correct_*/*50, with Wilson 95% binomial confidence intervals.
- **Coverage** = *n*_sure_*/*50, the fraction of questions where the agent did not invoke the refusal option.
- **Precision** = *n*_correct_*/n*_sure_, accuracy among questions the agent attempted.
- **Per-capsule accuracy**: pooled across the 1–4 questions per capsule.
- **Pairwise agreement on correctness**: fraction of questions on which two agents both succeed or both fail.

### 3.5 Data and code availability

Full per-question prediction CSVs (`combined.csv`), grading JSONs (`combined_graded.json`), and grading reports (`combined_grading_report.md`) are provided under `results/` in the `codex_agent`, `claude_code_agent`, and `codex_agent_5.5` subdirectories. Analysis scripts (`agent_results.py`, `make_figures.py`) and the per-question correctness matrix (`per_question_correctness.csv`) are provided alongside this manuscript.

## 4 Results

### 4.1 Overall accuracy

Table 2 and Figure 1 summarize overall performance. The two offline configurations achieve 84.0% (Opus 4.7) and 88.0% (GPT-5.4); the web-enabled GPT-5.5 configuration achieves 98.0%. Because each configuration was run once, we interpret the differences as an observed benchmark comparison rather than a replicate estimate of model variance. Coverage is high in all three configurations (92– 100%); the agents rarely invoked the refusal option.

**Table 2:** Overall results on BixBench-Verified-50 (*n* = 50). *n*_correct_: questions answered correctly; *n*_sure_: questions on which the agent did not return the refusal option; Coverage = *n*_sure_*/n*; Precision = *n*_correct_*/n*_sure_. 95% CI is Wilson binomial.

| Configuration | Accuracy | 95% CI | $n_{\text{correct}}/50$ | Coverage | Precision |
| --- | --- | --- | --- | --- | --- |
| GPT-5.4 (no web) | 88.0% | [76.2, 94.4] | 44 | 94.0% | 93.6% |
| Opus 4.7 (no web) | 84.0% | [71.5, 91.7] | 42 | 92.0% | 91.3% |
| GPT-5.5 (web) | **98.0%** | [89.5, 99.6] | **49** | 100.0% | 98.0% |

**Table 3:** Incorrect-answer summary. The web-enabled GPT-5.5 configuration has one official miss, `bix-16-q1`; review of the trajectory showed that the computation was correct for the raw `CRISPRGeneEffect.csv` sign convention, while the reference answer depends on interpreting essentiality with the opposite sign.

| Configuration | Official errors | Accuracy | Dominant error mode |
| --- | --- | --- | --- |
| GPT-5.4 (no web) | 6 | 88.0% | Missing offline pathway or phylogenetics resources, plus the shared essentiality sign-convention question. |
| Opus 4.7 (no web) | 8 | 84.0% | Mixture of numerical/preprocessing disagreements, pathway analysis errors, and the shared essentiality sign-convention question. |
| GPT-5.5 (web) | 1 | 98.0% | Only `bix-16-q1`; no clear computational failure after trajectory review. |

**Figure 1:**
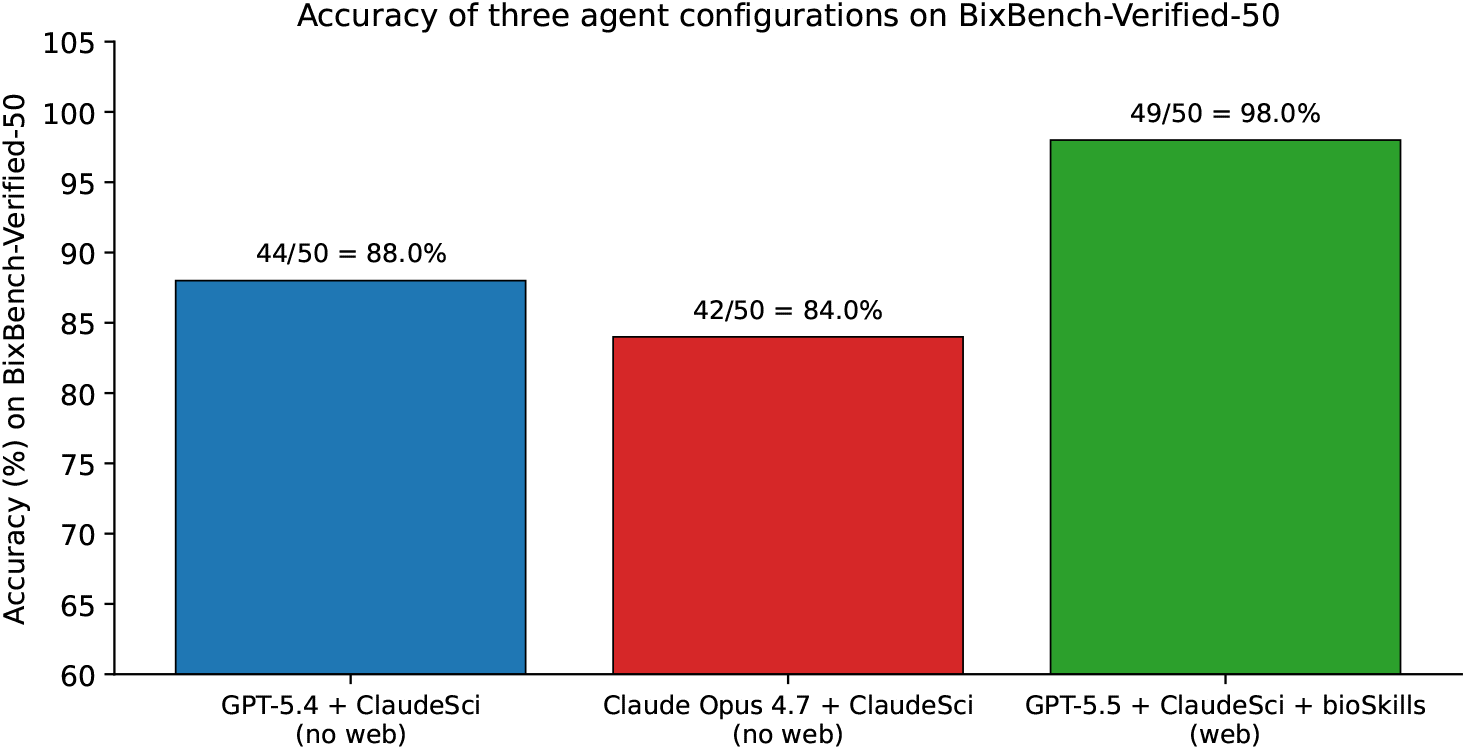
Accuracy of three frontier-model agent configurations on BixBench-Verified-50. Bars show one complete benchmark run per configuration; no run-level error bars are shown. GPT-5.5 with the bioSkills overlay and web access exceeds both offline configurations.

### 4.2 Comparison to published results

Figure 2 places these results in the context of previously reported BixBench-Verified-50 numbers. Our weaker offline configuration (Opus 4.7 + ClaudeSci, 84.0%) substantially exceeds the vanilla Claude Code (Opus 4.6) baseline reported by Phylo (65.3%) [2], although the model version and skill availability are both different. Our GPT-5.4 (no web) configuration at 88.0% is comparable to Biomni Lab’s reported 88.7% and within the band of K-Dense Web (90.0%) and SciAgent-Skills (92.0%). The GPT-5.5 (web) configuration sets a new high water mark for BixBench-Verified-50 at 98.0%, exceeding the previous best (92.0%) by 6 percentage points.

**Figure 2:**
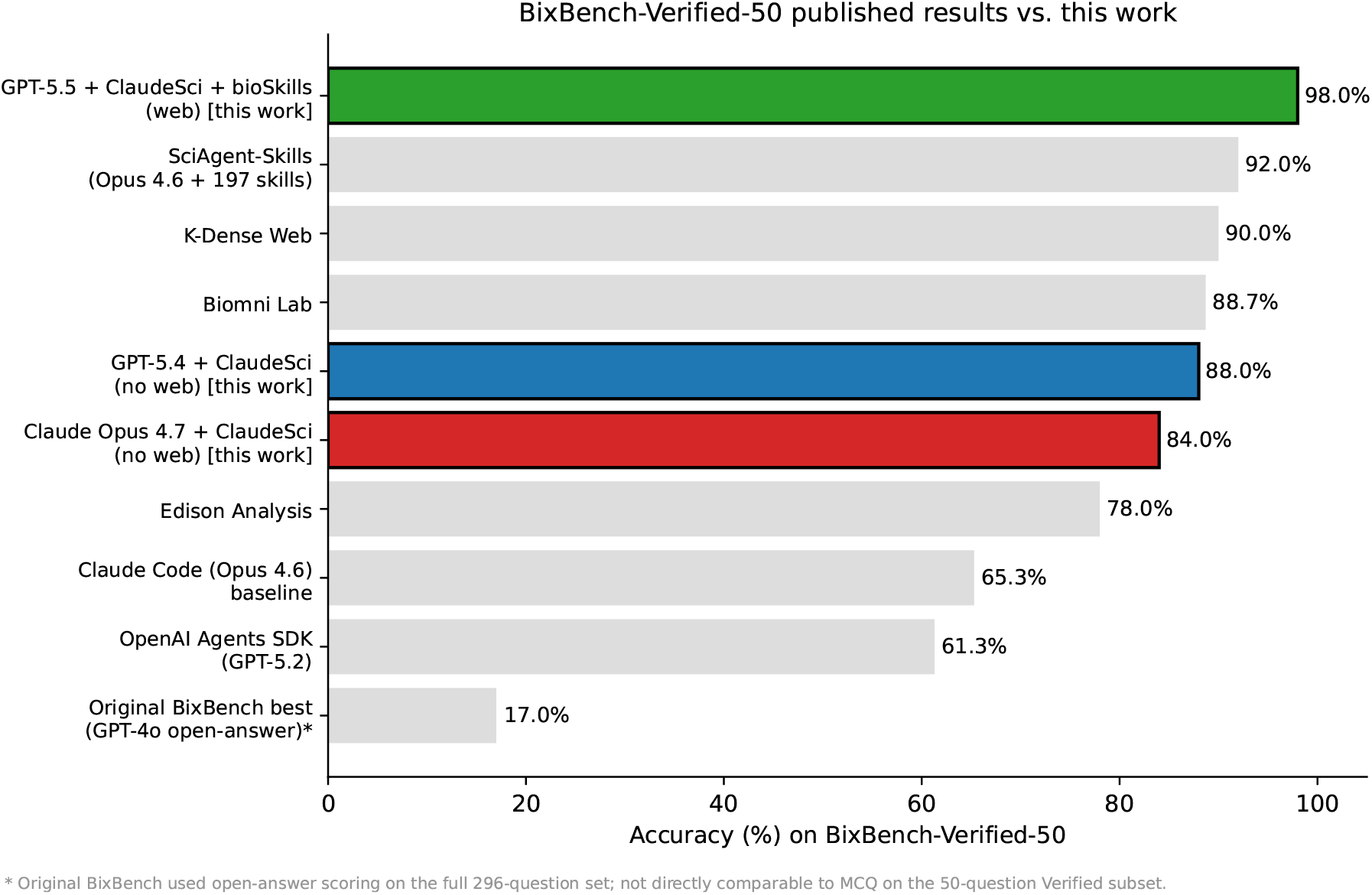
Reported accuracy on BixBench-Verified-50 across published agents and the three configurations evaluated in this work (highlighted). Numbers for prior systems were taken from the Phylo blog [2], the K-Dense Web blog [7], and the SciAgent-Skills repository [8]. The original BixBench frontier-model number is shown for context but used a different scoring protocol on the full 296-question set.

### 4.3 Per-capsule and per-category breakdown

Figure 3 shows per-capsule accuracy for all three configurations. 24 of 32 capsules are answered perfectly by all three agents (3 in green for every column). Difficulty concentrates in eight capsules, almost all of which are KEGG/pathway-enrichment problems (`bix-26`, `bix-32`, `bix-53`), phylogenetics problems requiring PhyKIT or BUSCO HMM databases (`bix-11`, `bix-12`, `bix-34`, `bix-38`), or expression/essentiality questions (`bix-16`). The web-enabled configuration recovers all but one of these (`bix-16`, discussed below).

**Figure 3:**
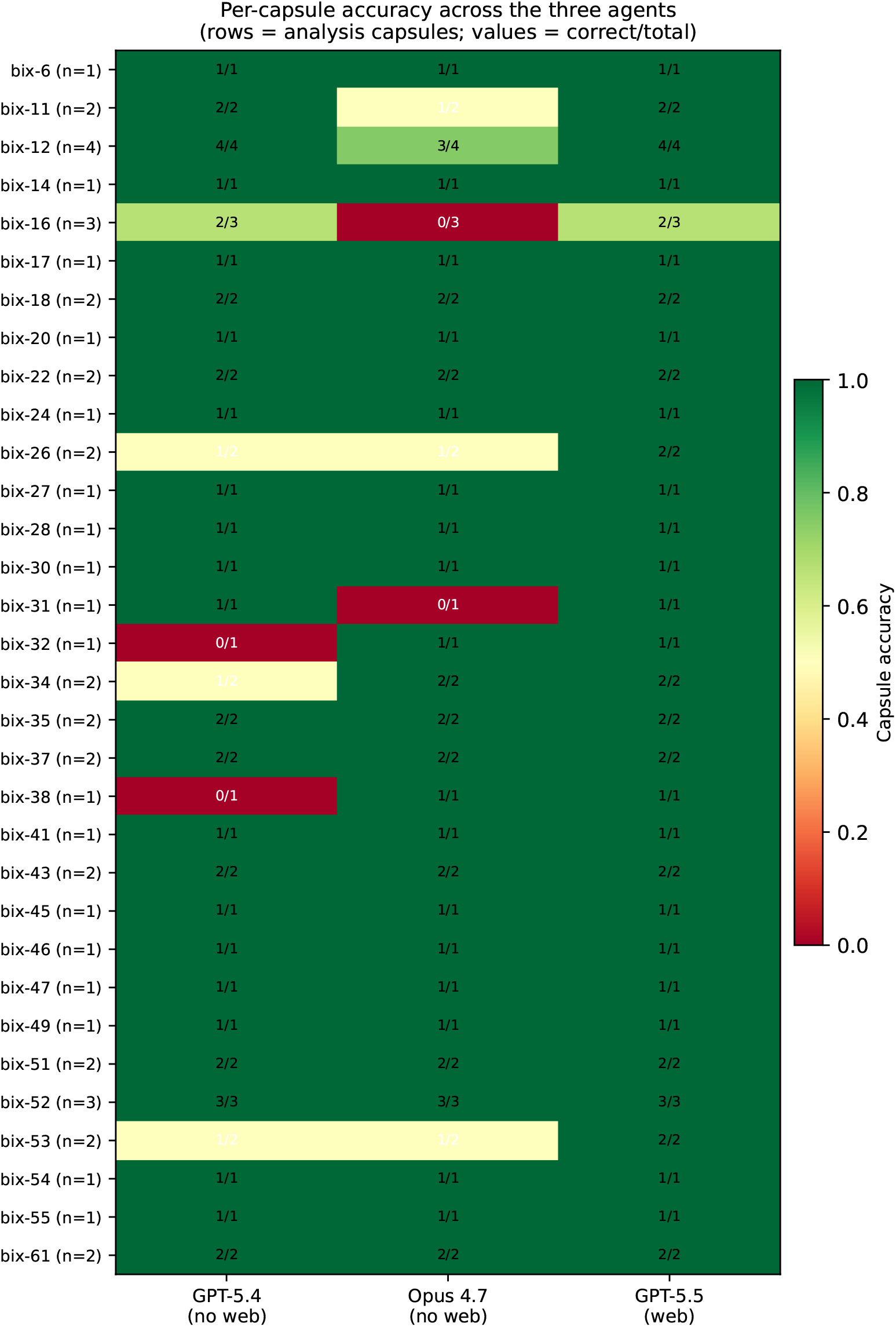
Per-capsule accuracy across the three agents. Green: capsule fully solved; red: all questions wrong; yellow: partial. Numbers in cells are correct/total questions for that capsule. Most failures cluster in capsules requiring external pathway annotations (KEGG, WikiPathways) or BUSCO/phylogenetics resources.

Figure 4 aggregates the same data into eight question categories. The web-enabled configuration is at 100% in every category except CRISPR/essentiality, where the residual error is the `bix-16-q1` failure shared by all three agents. The largest gain from web access + bioSkills appears in the differential-expression / pathway-enrichment family (12 questions), where it converts a 75% score (offline) into 100%. Phylogenetics/comparative-genomics is the second-largest beneficiary, recovering the BUSCO-HMM and PhyKIT failures that the offline agents could not resolve from the local capsule alone.

**Figure 4:**
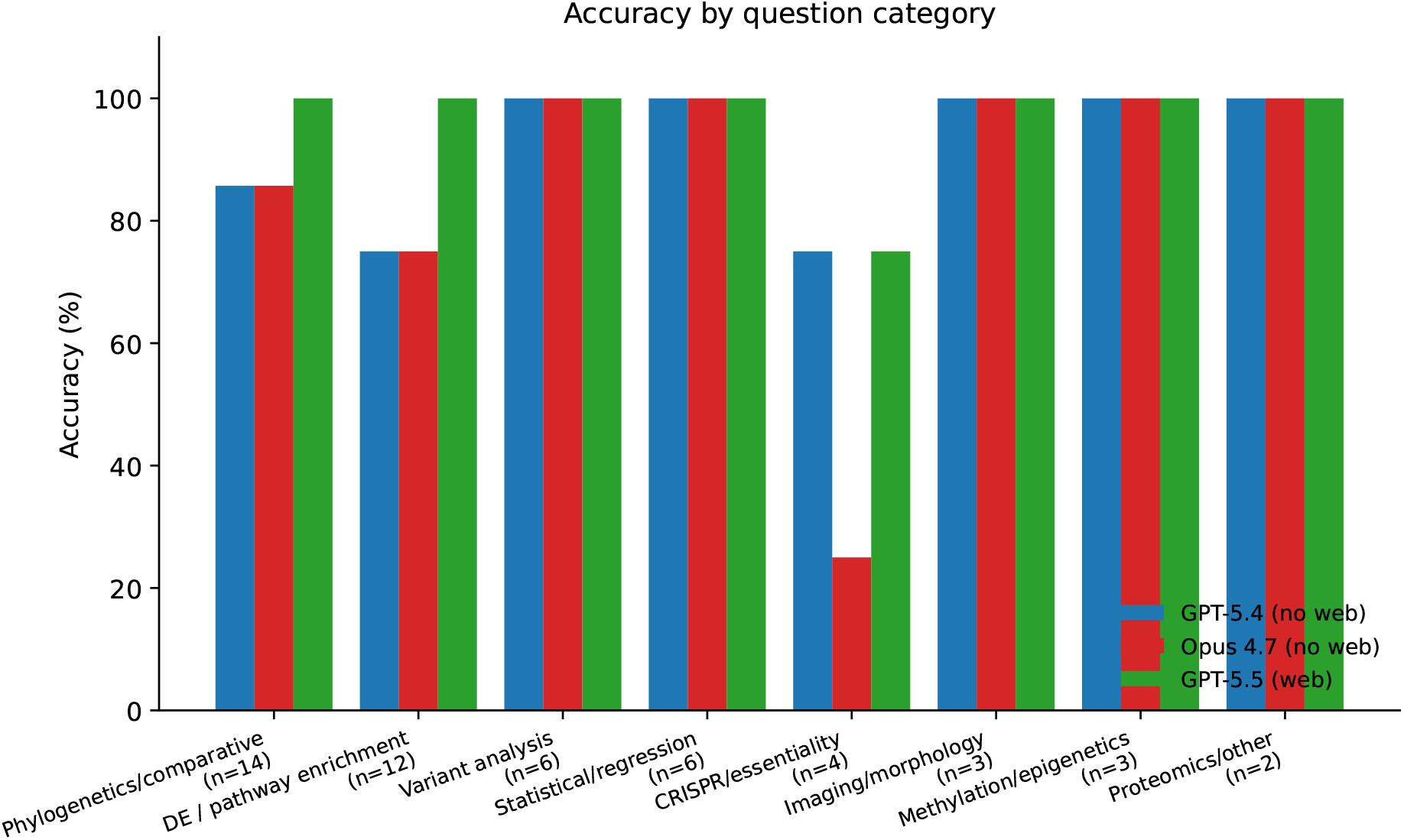
Accuracy by question category (capsules grouped by primary analytical task). Web-enabled GPT-5.5 reaches 100% in every category except CRISPR/essentiality, where the single residual error is shared by all three agents.

### 4.4 Agreement and shared errors

The three agents disagree primarily through the offline pair making different mistakes (Figure 5). Pairwise agreement on correctness is 76% between the two offline agents (GPT-5.4 vs. Opus 4.7), 90% between GPT-5.4 (no web) and GPT-5.5 (web), and 86% between Opus 4.7 (no web) and GPT-5.5 (web). Strikingly, only one question (`bix-16-q1`) is missed by all three agents.

**Figure 5:**
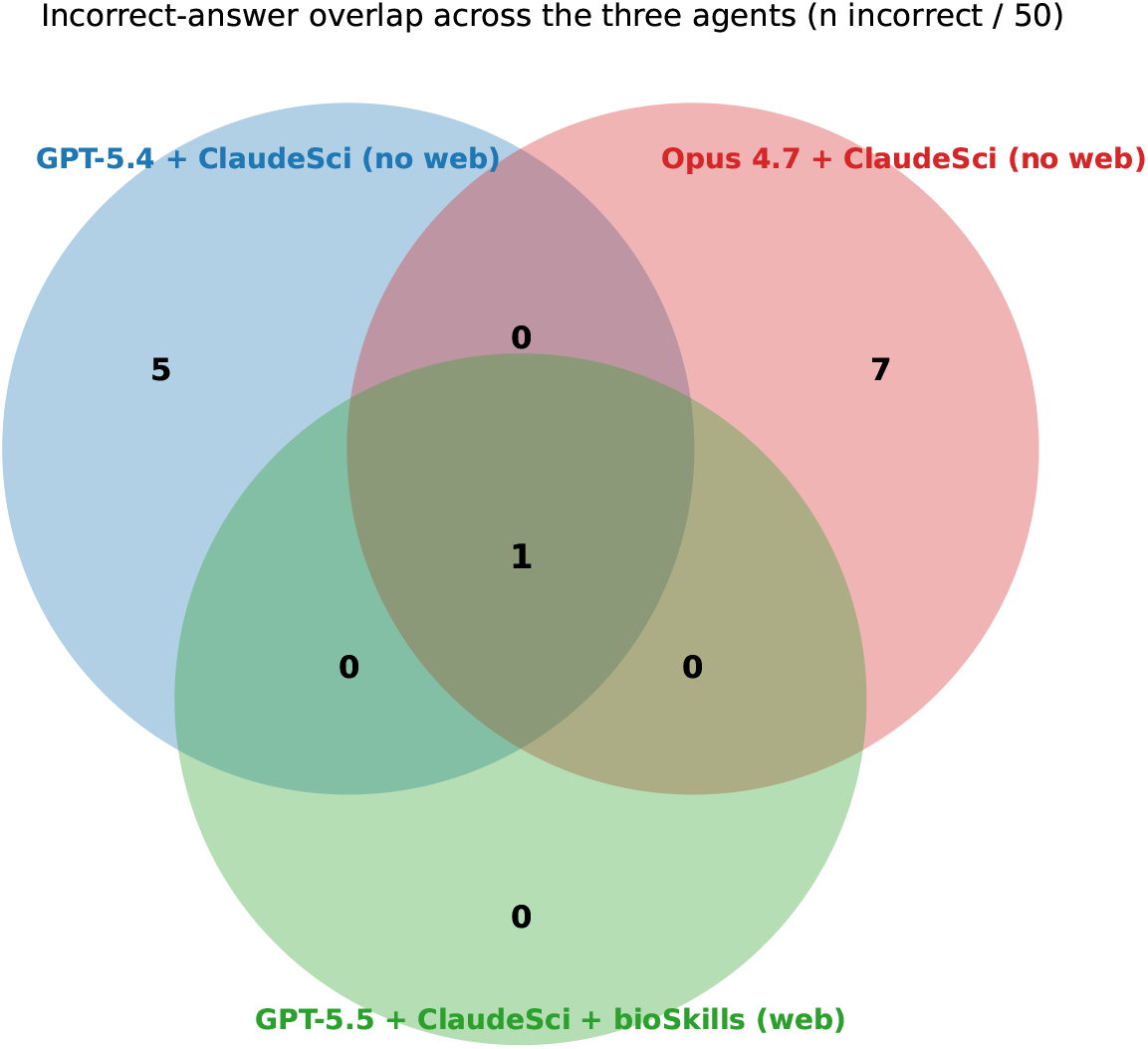
Set diagram of incorrect-answer overlap. The two offline configurations make 5 (GPT-5.4) and 7 (Opus 4.7) errors that are unique to one of them; only one question is missed by all three configurations.

### 4.5 Qualitative error analysis

We examined every incorrect answer in detail. Errors fall into three categories.

i. **Missing offline annotations**. The clearest pattern in the offline configurations is failure on questions that require external annotation databases the agent does not have locally. GPT-5.4 (no web) explicitly notes this in its agent summaries: “*KEGG*.*db and PA14 OrgDb packages are not installed, and* `clusterProfiler::enrichKEGG` *fails offline when trying to fetch* `pau` *pathway mappings*” (`bix-26-q3`); “*the capsule only contains three DESeq2 result files and no KEGG pathway mapping*” (`bix-32-q2`); “*no installed gseapy, no local WikiPathways_2019_Mouse library, and no local mouse annotation package to map the Ensembl mouse gene IDs to pathway-library gene symbols*” (`bix-53-q5`). In each of these cases the web-enabled GPT-5.5 successfully fetches the necessary library or annotation and answers correctly. Two phylogenetics questions (`bix-38-q1`, `bix-34-q2`) fail in the offline GPT-5.4 configuration with similar root cause (no installed PhyKIT, no BUSCO HMM database for the requested gene).
ii. **Off-by-one numerical disagreement**. Several offline errors involve small numerical disagreements with the gold answer, often within a factor of 2 of the correct value. The Opus 4.7 errors on `bix-31-q2` (FAM138A log_2_FC) and `bix-12-q6` (Mann–Whitney *U* statistic) are of this flavor; the chosen distractor is plausible but reflects a different shrinkage method or a different preprocessing step than the gold notebook used.
iii. **Persistent shared error:** `bix-16-q1`. The question asks for the gene with the strongest negative Spearman correlation between expression and essentiality. All three agents used the same basic formulation: align expression and `CRISPRGeneEffect.csv` by shared cell-line model, compute gene-wise Spearman correlations, and select the most negative coefficient. Under the raw sign convention in the distributed CRISPR gene-effect file, this procedure yields CCND1 (*ρ* = −0.629), with KLF5 at *ρ* = −0.602 in the GPT-5.5 run. The reference answer is CDKN1A. Reversing the sign of the gene-effect scores before correlation changes which direction corresponds to increasing essentiality and is consistent with the reference answer. We therefore count `bix-16-q1` as an official benchmark miss, but interpret it as a question-wording and sign-convention ambiguity rather than evidence that the agent failed to perform the requested statistical analysis.

## 5 Discussion

### 5.1 Are SOTA agents ready for bioinformatics?

The headline answer from this work is that, for the kind of short, well-specified analytical question represented in BixBench-Verified-50, frontier agents with appropriate scientific skills are now within one question of perfect performance. The single remaining GPT-5.5 miss appears to be a sign-convention ambiguity rather than a failure to load, clean, analyze, or statistically interpret the data. This is a substantial shift from the picture in the original BixBench paper a year ago, and it calls for a narrower conclusion than either optimism or skepticism alone would support: these agents are already highly capable on bounded, well-specified bioinformatics tasks, but that does not mean they can yet replace expert oversight on open-ended research projects.

The 88%-then-98% pattern across our two GPT configurations is particularly informative because the no-web GPT-5.4 configuration already performs in the same range as previously reported leading systems. This suggests that a competent base model plus a curated scientific-skill library is sufficient for many routine benchmark tasks where the required data are local and the expected analysis is well defined. Web access changes the picture mainly by filling gaps in offline annotation databases and software packages (KEGG, WikiPathways, PhyKIT, BUSCO HMM models, mouse OrgDb), a narrow but common class of problems in practical bioinformatics.

### 5.2 Where the easy gains came from

Our category-level breakdown (Figure 4) makes the locus of improvement explicit. Variant analysis, statistical/regression, imaging/morphology, methylation, and proteomics are at 100% for all three configurations: these are tasks where everything the agent needs is in the capsule. Phylogenetics and DE/pathway-enrichment are the categories where web access matters, because both rely on external annotation resources or specialized software (PhyKIT, BUSCO, clusterProfiler, gseapy with Mouse pathway libraries). Within these two categories, the “hard” questions are uniformly resolved by GPT-5.5 (web) once the agent can install the required package or fetch the required pathway file. This is consistent with the SciAgent-Skills report [8], which observed a +26.7-point lift on BixBench-Verified-50 simply by equipping Claude Code (Opus 4.6) with 197 bioinformatics skills, no fine-tuning required.

### 5.3 Limitations of BixBench-Verified-50 as an “agent readiness” test

While BixBench-Verified-50 is the cleanest publicly available agent benchmark for bioinformatics, it has structural features that make it easier than typical research analysis. (a) Questions are multiple-choice with at most five options; the prior on a random guess is 25% rather than effectively zero for an open answer. (b) Each question is associated with one curated capsule that contains the relevant input data; the agent does not have to discover the data or determine its provenance. (c) The questions are derived from published, peer-reviewed analytical workflows, so a correct procedure provably exists; this is not the situation in exploratory analysis. (d) The “Insufficient information” refusal option is itself a hint to a calibrated agent that the question is answerable. (e) The 50 questions are short — typically under a few hundred LOC of code on the agent side — whereas a real dissertation chapter analysis might be tens of thousands of lines.

The mismatch between BixBench-style questions and the real day-to-day work of a computational biologist is the central caveat to our conclusion. Phylo’s recent BiomniBench framework [2] attempts to address this by scoring agents on the entire analytical trace rather than just the final number; we view that direction as strictly complementary to the work here. Saturating BixBench-Verified-50 is necessary but probably not sufficient for “agent readiness”.

### 5.4 Implications for skill libraries

Our results suggest that the most cost-effective way to upgrade a generic frontier-model coding agent for bioinformatics is to attach a curated skill library, not to build a bespoke multi-agent architecture. The comparison is not a perfect ablation, because model versions differ across public leaderboard entries, but the pattern is consistent across our runs and external reports: agents with explicit scientific skills perform far better than vanilla coding-agent baselines on the same verified benchmark [2, 7, 8]. Stacking a second skill library (bioSkills) plus web access on top of a stronger base model yields the highest score we have seen reported. Skill libraries also have the appealing property that they are inspectable, version-controlled markdown [12], and therefore reviewable by domain experts in a way that a fine-tuned weights-only model is not.

### 5.5 What’s still missing

Three things would change our headline conclusion. First, we did not evaluate end-to-end *open-answer* performance on these capsules; the original BixBench numbers were dramatically lower in open-answer mode, and skill libraries may not close that gap as completely as they close the MCQ gap. Second, we did not measure trace quality: a 98% accuracy agent that arrives at correct answers via incorrect intermediate steps would still mislead a human collaborator. Third, the questions in BixBench-Verified-50 are designed around 1–3 day analyses; nothing in this benchmark validates that an agent can autonomously sustain multi-week analytical projects with shifting hypotheses, which is arguably what real “agent readiness” for biology research would require.

## 6 Conclusion

Frontier-model agents equipped with curated scientific skills are markedly more capable on bioinformatics tasks than they were a year ago. On BixBench-Verified-50 our offline configurations (84% and 88%) already match the previously reported leaderboard band, and our web-enabled GPT-5.5 + Claude Scientific Skills + bioSkills configuration sets a new high (98%, 49/50), with only one official miss and no clear computational failure after trajectory review. The locus of improvement is concentrated in tasks that require external annotation databases or specialized software: web access is not magic, it is a targeted fix for offline gaps. Skill libraries are an unusually cost-effective lever, contributing substantial headline accuracy while remaining inspectable by domain experts. We caution that BixBench-Verified-50 measures a particular, narrow slice of bioinformatics research; near-saturation is necessary but not sufficient evidence that frontier agents can replace experienced computational biologists. Trace-level evaluations [2, 3], open-answer benchmarks, and longer analytical projects are the natural next tests.

## Supporting information

evaluation results and scripts

## Data and Code Availability

Per-agent prediction CSVs (`combined.csv`), grading JSONs (`combined_graded.json`), grading reports (`combined_grading_report.md`), the per-question correctness matrix, figure-generation code, and per-capsule analysis script are bundled with this manuscript. BixBench-Verified-50 is available at https://huggingface.co/datasets/phylobio/BixBench-Verified-50.

## Acknowledgments

We thank the FutureHouse and ScienceMachine teams for releasing BixBench, the Phylo team for curating BixBench-Verified-50, and the maintainers of Claude Scientific Skills (K-Dense AI) and bioSkills (GPTomics) for releasing their skill libraries under permissive licences.

## Footnotes

1 Capsule `bix-16` appears with three questions in our subset and the dataset description on Hugging Face reports 33 capsules; the difference of one is consistent with our local mapping of question UUIDs to capsule UUIDs.

