## Supplementary material for "Skill-Augmented Frontier Agents Nearly Saturate BixBench-Verified-50": evaluation results and scripts: agent_results.py

```
"""
Per-agent per-question correctness on BixBench-Verified-50.
Extracted from each agent's combined_grading_report.md.
"""

# Order from codex_agent's correct (44) + incorrect (6) lists, total 50.
ALL_QUESTIONS = [
    "bix-6-q4","bix-61-q5","bix-53-q2","bix-27-q5","bix-54-q7","bix-51-q2","bix-12-q5",
    "bix-30-q3","bix-31-q2","bix-47-q3","bix-12-q2","bix-35-q2","bix-34-q5","bix-11-q1",
    "bix-35-q1","bix-16-q3","bix-14-q1","bix-20-q3","bix-55-q1","bix-12-q4","bix-41-q5",
    "bix-43-q4","bix-37-q1","bix-52-q7","bix-22-q1","bix-12-q6","bix-24-q2","bix-46-q4",
    "bix-37-q4","bix-26-q5","bix-45-q1","bix-18-q3","bix-11-q2","bix-16-q4","bix-22-q4",
    "bix-61-q2","bix-51-q8","bix-17-q2","bix-43-q2","bix-18-q1","bix-28-q3","bix-52-q2",
    "bix-52-q6","bix-49-q4",
    "bix-26-q3","bix-32-q2","bix-53-q5","bix-16-q1","bix-38-q1","bix-34-q2",
]
assert len(ALL_QUESTIONS) == 50

# Incorrect sets per agent, derived from each combined_grading_report.md
INCORRECT = {
    "codex54":   {"bix-26-q3","bix-32-q2","bix-53-q5","bix-16-q1","bix-38-q1","bix-34-q2"},
    "claude47":  {"bix-53-q2","bix-31-q2","bix-16-q3","bix-11-q1","bix-16-q1","bix-12-q6","bix-26-q5","bix-16-q4"},
    "codex55":   {"bix-16-q1"},
}

AGENT_LABEL = {
    "codex54":  "GPT-5.4 + Claude-Sci (no web)",
    "claude47": "Claude Opus 4.7 + Claude-Sci (no web)",
    "codex55":  "GPT-5.5 + Claude-Sci + bioSkills (web)",
}

# Capsule -> source paper mapping; here we just track capsules per question family.
import re
def capsule_id(qid):
    m = re.match(r"^(bix-\d+)-q\d+$", qid)
    return m.group(1) if m else qid

CAPSULE_OF = {q: capsule_id(q) for q in ALL_QUESTIONS}
N_CAPSULES = len(set(CAPSULE_OF.values()))
print(f"Total questions: {len(ALL_QUESTIONS)}")
print(f"Distinct capsules: {N_CAPSULES}")

# Per-agent accuracy
print("\n=== Accuracy ===")
for a, inc in INCORRECT.items():
    n_correct = len(ALL_QUESTIONS) - len(inc)
    print(f"{a}: {n_correct}/50 = {100*n_correct/50:.1f}% - {AGENT_LABEL[a]}")

# Wilson 95% CI
from math import sqrt
def wilson(n_correct, n, z=1.96):
    p = n_correct / n
    denom = 1 + z*z/n
    center = (p + z*z/(2*n)) / denom
    half = (z * sqrt((p*(1-p)/n) + z*z/(4*n*n))) / denom
    return (center-half, center+half)
print("\n=== Wilson 95% CI ===")
for a, inc in INCORRECT.items():
    n_correct = 50 - len(inc)
    lo, hi = wilson(n_correct, 50)
    print(f"{a}: [{100*lo:.1f}%, {100*hi:.1f}%]")

# Per-question correctness matrix
rows = []
for q in ALL_QUESTIONS:
    cap = CAPSULE_OF[q]
    c54 = 0 if q in INCORRECT["codex54"]  else 1
    c47 = 0 if q in INCORRECT["claude47"] else 1
    c55 = 0 if q in INCORRECT["codex55"]  else 1
    rows.append((q, cap, c54, c47, c55))

# Group by capsule
from collections import defaultdict
cap_correct = defaultdict(lambda: {"codex54":[], "claude47":[], "codex55":[]})
for q,cap,c54,c47,c55 in rows:
    cap_correct[cap]["codex54"].append(c54)
    cap_correct[cap]["claude47"].append(c47)
    cap_correct[cap]["codex55"].append(c55)

print("\n=== Capsule-level accuracy ===")
print(f"{'capsule':<10}{'n_q':<5}{'codex54':<10}{'claude47':<10}{'codex55':<10}")
for cap in sorted(cap_correct, key=lambda x:int(x.split('-')[1])):
    nq = len(cap_correct[cap]["codex54"])
    s54 = sum(cap_correct[cap]["codex54"])
    s47 = sum(cap_correct[cap]["claude47"])
    s55 = sum(cap_correct[cap]["codex55"])
    print(f"{cap:<10}{nq:<5}{s54}/{nq:<8}{s47}/{nq:<8}{s55}/{nq}")

# Pairwise agreement on correctness
def agree(set_a, set_b):
    """Both right or both wrong on the same question."""
    same = 0
    for q in ALL_QUESTIONS:
        a = q not in set_a
        b = q not in set_b
        if a == b: same += 1
    return same / len(ALL_QUESTIONS)

print("\n=== Pairwise correctness agreement ===")
keys = list(INCORRECT.keys())
for i in range(len(keys)):
    for j in range(i+1, len(keys)):
        ag = agree(INCORRECT[keys[i]], INCORRECT[keys[j]])
        print(f"{keys[i]} vs {keys[j]}: {100*ag:.1f}%")

# Questions hard for ALL three (intersection of incorrect sets)
shared_hard = INCORRECT["codex54"] & INCORRECT["claude47"] & INCORRECT["codex55"]
print(f"\nQuestions ALL agents got wrong: {sorted(shared_hard)}")

# Save tidy CSV
import csv, os
out_dir = os.path.dirname(os.path.abspath(__file__))
with open(os.path.join(out_dir, "per_question_correctness.csv"), "w", newline="") as fh:
    w = csv.writer(fh)
    w.writerow(["question_id","capsule","codex54_correct","claude47_correct","codex55_correct"])
    for r in rows:
        w.writerow(r)
print("\nWrote per_question_correctness.csv")
```
