## Supplementary material for "Skill-Augmented Frontier Agents Nearly Saturate BixBench-Verified-50": evaluation results and scripts: make_figures.py

```
"""
Build publication-quality figures for the BixBench-Verified-50 paper.

Run from this directory:

    pip install matplotlib numpy
    python make_figures.py

Reads per_question_correctness.csv and writes the five paper figures
into ./figures/ as both PDF (for LaTeX include) and PNG (for previews).

Figures:
  fig1_accuracy.pdf       - Accuracy of the 3 agents from a single run
  fig2_leaderboard.pdf    - Our agents vs published BixBench-Verified-50 results
  fig3_capsule_heatmap.pdf - Per-capsule correctness across the 3 agents
  fig4_venn.pdf           - Overlap of incorrect questions across the 3 agents
  fig5_categories.pdf     - Accuracy by question category
"""
import os, csv, math
import matplotlib
matplotlib.use("Agg")
import matplotlib.pyplot as plt
from matplotlib.patches import Circle
import numpy as np

OUT = os.path.dirname(os.path.abspath(__file__))
FIG = os.path.join(OUT, "figures")
os.makedirs(FIG, exist_ok=True)

COLORS = {
    "codex54":  "#1f77b4",
    "claude47": "#d62728",
    "codex55":  "#2ca02c",
}

AGENTS = [
    ("codex54",  "GPT-5.4 + ClaudeSci\n(no web)",         44, 50, COLORS["codex54"]),
    ("claude47", "Claude Opus 4.7 + ClaudeSci\n(no web)", 42, 50, COLORS["claude47"]),
    ("codex55",  "GPT-5.5 + ClaudeSci + bioSkills\n(web)",49, 50, COLORS["codex55"]),
]

def wilson(k, n, z=1.96):
    p = k/n
    denom = 1 + z*z/n
    center = (p + z*z/(2*n)) / denom
    half = (z * math.sqrt((p*(1-p)/n) + z*z/(4*n*n))) / denom
    return center, max(0, center-half), min(1, center+half)

# ---------- Figure 1: our three agents ----------
fig, ax = plt.subplots(figsize=(7.5, 4.0))
labels = [a[1] for a in AGENTS]
means  = [a[2]/a[3] for a in AGENTS]
colors = [a[4] for a in AGENTS]
xs = np.arange(len(AGENTS))
ax.bar(xs, [m*100 for m in means],
       color=colors, edgecolor="black", linewidth=0.6)
for x, m, a in zip(xs, means, AGENTS):
    ax.text(x, m*100 + 1.0, f"{a[2]}/{a[3]} = {m*100:.1f}%",
            ha="center", va="bottom", fontsize=9)
ax.set_xticks(xs)
ax.set_xticklabels(labels, fontsize=9)
ax.set_ylabel("Accuracy (%) on BixBench-Verified-50")
ax.set_ylim(60, 105)
ax.spines["top"].set_visible(False); ax.spines["right"].set_visible(False)
ax.set_title("Accuracy of three agent configurations on BixBench-Verified-50",
             fontsize=11)
plt.tight_layout()
plt.savefig(os.path.join(FIG, "fig1_accuracy.pdf"))
plt.savefig(os.path.join(FIG, "fig1_accuracy.png"), dpi=180)
plt.close()

# ---------- Figure 2: leaderboard context ----------
LEADER = [
    ("Original BixBench best\n(GPT-4o open-answer)*",          17.0, "lit",  "lightgray"),
    ("Claude Code (Opus 4.6)\nbaseline",                       65.3, "lit",  "lightgray"),
    ("OpenAI Agents SDK\n(GPT-5.2)",                           61.3, "lit",  "lightgray"),
    ("Edison Analysis",                                        78.0, "lit",  "lightgray"),
    ("Biomni Lab",                                             88.7, "lit",  "lightgray"),
    ("K-Dense Web",                                            90.0, "lit",  "lightgray"),
    ("SciAgent-Skills\n(Opus 4.6 + 197 skills)",               92.0, "lit",  "lightgray"),
    ("Claude Opus 4.7 + ClaudeSci\n(no web) [this work]",      84.0, "ours", COLORS["claude47"]),
    ("GPT-5.4 + ClaudeSci\n(no web) [this work]",              88.0, "ours", COLORS["codex54"]),
    ("GPT-5.5 + ClaudeSci + bioSkills\n(web) [this work]",     98.0, "ours", COLORS["codex55"]),
]
LEADER_sorted = sorted(LEADER, key=lambda x: x[1])
fig, ax = plt.subplots(figsize=(8.5, 5.5))
ys = np.arange(len(LEADER_sorted))
vals = [x[1] for x in LEADER_sorted]
cols = [x[3] for x in LEADER_sorted]
edges = ["black" if x[2] == "ours" else "none" for x in LEADER_sorted]
ax.barh(ys, vals, color=cols, edgecolor=edges, linewidth=1.4)
for y, x in zip(ys, LEADER_sorted):
    ax.text(x[1] + 0.7, y, f"{x[1]:.1f}%", va="center", fontsize=9)
ax.set_yticks(ys)
ax.set_yticklabels([x[0] for x in LEADER_sorted], fontsize=8.5)
ax.set_xlabel("Accuracy (%) on BixBench-Verified-50")
ax.set_xlim(0, 105)
ax.set_title("BixBench-Verified-50 published results vs. this work", fontsize=11)
ax.spines["top"].set_visible(False); ax.spines["right"].set_visible(False)
note = ("* Original BixBench used open-answer scoring on the full 296-question set; "
        "not directly comparable to MCQ on the 50-question Verified subset.")
fig.text(0.02, 0.005, note, fontsize=7, color="gray")
plt.tight_layout(rect=[0, 0.02, 1, 1])
plt.savefig(os.path.join(FIG, "fig2_leaderboard.pdf"))
plt.savefig(os.path.join(FIG, "fig2_leaderboard.png"), dpi=180)
plt.close()

# ---------- Figure 3: per-capsule heatmap ----------
QC = []
with open(os.path.join(OUT, "per_question_correctness.csv")) as fh:
    r = csv.DictReader(fh)
    for row in r:
        QC.append(row)

def cap_index(c):
    return int(c.split("-")[1])
caps = sorted(set(q["capsule"] for q in QC), key=cap_index)
mat = np.zeros((len(caps), 3))
nq_per_cap = []
for i, cap in enumerate(caps):
    rows = [q for q in QC if q["capsule"] == cap]
    nq_per_cap.append(len(rows))
    mat[i, 0] = np.mean([int(r["codex54_correct"])  for r in rows])
    mat[i, 1] = np.mean([int(r["claude47_correct"]) for r in rows])
    mat[i, 2] = np.mean([int(r["codex55_correct"])  for r in rows])

fig, ax = plt.subplots(figsize=(6.5, 9.5))
cmap = plt.cm.RdYlGn
im = ax.imshow(mat, cmap=cmap, vmin=0, vmax=1, aspect="auto")
ax.set_xticks([0, 1, 2])
ax.set_xticklabels(["GPT-5.4\n(no web)", "Opus 4.7\n(no web)", "GPT-5.5\n(web)"], fontsize=9)
ax.set_yticks(range(len(caps)))
ax.set_yticklabels([f"{c} (n={nq})" for c, nq in zip(caps, nq_per_cap)], fontsize=8)
for i in range(len(caps)):
    for j in range(3):
        v = mat[i, j]
        txt = f"{int(round(v*nq_per_cap[i]))}/{nq_per_cap[i]}"
        ax.text(j, i, txt, ha="center", va="center", fontsize=7,
                color="black" if v > 0.5 else "white")
cbar = plt.colorbar(im, ax=ax, fraction=0.04, pad=0.02)
cbar.set_label("Capsule accuracy", fontsize=9)
ax.set_title("Per-capsule accuracy across the three agents\n"
             "(rows = analysis capsules; values = correct/total)", fontsize=10)
plt.tight_layout()
plt.savefig(os.path.join(FIG, "fig3_capsule_heatmap.pdf"))
plt.savefig(os.path.join(FIG, "fig3_capsule_heatmap.png"), dpi=180)
plt.close()

# ---------- Figure 4: Venn-like diagram of incorrect questions ----------
INC = {
    "codex54":  {"bix-26-q3","bix-32-q2","bix-53-q5","bix-16-q1","bix-38-q1","bix-34-q2"},
    "claude47": {"bix-53-q2","bix-31-q2","bix-16-q3","bix-11-q1","bix-16-q1","bix-12-q6","bix-26-q5","bix-16-q4"},
    "codex55":  {"bix-16-q1"},
}
A, B, C = INC["codex54"], INC["claude47"], INC["codex55"]
abc = A & B & C
counts = {
    "A_only": len(A - B - C),
    "B_only": len(B - A - C),
    "C_only": len(C - A - B),
    "AB":     len((A & B) - abc),
    "AC":     len((A & C) - abc),
    "BC":     len((B & C) - abc),
    "ABC":    len(abc),
}
fig, ax = plt.subplots(figsize=(7.0, 6.0))
ax.set_aspect("equal"); ax.axis("off")
ax.add_patch(Circle((-0.55,  0.30), 1.10, color=COLORS["codex54"],  alpha=0.35))
ax.add_patch(Circle(( 0.55,  0.30), 1.10, color=COLORS["claude47"], alpha=0.35))
ax.add_patch(Circle(( 0.00, -0.50), 1.10, color=COLORS["codex55"],  alpha=0.35))

def lab(x, y, t, fs=11):
    ax.text(x, y, t, ha="center", va="center", fontsize=fs, fontweight="bold")
lab(-1.15,  0.55, str(counts["A_only"]))
lab( 1.15,  0.55, str(counts["B_only"]))
lab( 0.00, -1.20, str(counts["C_only"]))
lab( 0.00,  0.65, str(counts["AB"]))
lab(-0.55, -0.40, str(counts["AC"]))
lab( 0.55, -0.40, str(counts["BC"]))
lab( 0.00, -0.05, str(counts["ABC"]), fs=12)

ax.text(-1.55,  1.20, "GPT-5.4 + ClaudeSci (no web)",
        color=COLORS["codex54"],  fontsize=10, fontweight="bold")
ax.text( 0.45,  1.20, "Opus 4.7 + ClaudeSci (no web)",
        color=COLORS["claude47"], fontsize=10, fontweight="bold")
ax.text(-0.85, -1.55, "GPT-5.5 + ClaudeSci + bioSkills (web)",
        color=COLORS["codex55"],  fontsize=10, fontweight="bold")
ax.set_xlim(-2.0, 2.2); ax.set_ylim(-1.8, 1.6)
ax.set_title("Incorrect-answer overlap across the three agents (n incorrect / 50)",
             fontsize=11)
plt.tight_layout()
plt.savefig(os.path.join(FIG, "fig4_venn.pdf"))
plt.savefig(os.path.join(FIG, "fig4_venn.png"), dpi=180)
plt.close()

# ---------- Figure 5: accuracy by category ----------
CATEGORY = {
    "bix-6":  "CRISPR/essentiality",
    "bix-11": "Phylogenetics/comparative",
    "bix-12": "Phylogenetics/comparative",
    "bix-14": "Variant analysis",
    "bix-16": "CRISPR/essentiality",
    "bix-17": "Variant analysis",
    "bix-18": "Imaging/morphology",
    "bix-20": "Variant analysis",
    "bix-22": "Statistical/regression",
    "bix-24": "DE / pathway enrichment",
    "bix-26": "DE / pathway enrichment",
    "bix-27": "Statistical/regression",
    "bix-28": "Phylogenetics/comparative",
    "bix-30": "DE / pathway enrichment",
    "bix-31": "DE / pathway enrichment",
    "bix-32": "DE / pathway enrichment",
    "bix-34": "Phylogenetics/comparative",
    "bix-35": "Phylogenetics/comparative",
    "bix-37": "Proteomics/other",
    "bix-38": "Phylogenetics/comparative",
    "bix-41": "Imaging/morphology",
    "bix-43": "DE / pathway enrichment",
    "bix-45": "Phylogenetics/comparative",
    "bix-46": "DE / pathway enrichment",
    "bix-47": "Variant analysis",
    "bix-49": "DE / pathway enrichment",
    "bix-51": "Statistical/regression",
    "bix-52": "Methylation/epigenetics",
    "bix-53": "DE / pathway enrichment",
    "bix-54": "Statistical/regression",
    "bix-55": "Phylogenetics/comparative",
    "bix-61": "Variant analysis",
}
cat_correct = {}
for q in QC:
    cat = CATEGORY.get(q["capsule"], "Other")
    d = cat_correct.setdefault(cat, {"codex54": [], "claude47": [], "codex55": [], "n": 0})
    d["codex54"].append(int(q["codex54_correct"]))
    d["claude47"].append(int(q["claude47_correct"]))
    d["codex55"].append(int(q["codex55_correct"]))
    d["n"] += 1

cats = sorted(cat_correct.keys(), key=lambda c: -cat_correct[c]["n"])
fig, ax = plt.subplots(figsize=(8.0, 5.0))
xs = np.arange(len(cats))
w = 0.27
for i, (k, col, lbl) in enumerate([
    ("codex54",  COLORS["codex54"],  "GPT-5.4 (no web)"),
    ("claude47", COLORS["claude47"], "Opus 4.7 (no web)"),
    ("codex55",  COLORS["codex55"],  "GPT-5.5 (web)"),
]):
    means = [100 * np.mean(cat_correct[c][k]) for c in cats]
    ax.bar(xs + (i - 1) * w, means, width=w, color=col, label=lbl)
ax.set_xticks(xs)
ax.set_xticklabels([f"{c}\n(n={cat_correct[c]['n']})" for c in cats],
                   rotation=20, ha="right", fontsize=8)
ax.set_ylabel("Accuracy (%)")
ax.set_ylim(0, 110)
ax.legend(loc="lower right", fontsize=8.5, frameon=False)
ax.spines["top"].set_visible(False); ax.spines["right"].set_visible(False)
ax.set_title("Accuracy by question category", fontsize=11)
plt.tight_layout()
plt.savefig(os.path.join(FIG, "fig5_categories.pdf"))
plt.savefig(os.path.join(FIG, "fig5_categories.png"), dpi=180)
plt.close()

print("All figures written to:", FIG)
for f in sorted(os.listdir(FIG)):
    print(" -", f)
```
